# Community diversity affects functionality and species sorting during propagation of a natural microbial community

**DOI:** 10.1101/2023.02.24.529839

**Authors:** Alanna Leale, Ben Auxier, Eddy J. Smid, Sijmen Schoustra

## Abstract

The influence of community diversity on community function has long been a central question in ecology. Particularly, the dynamics over time of this relationship as a function of levels of diversity remains unclear. Natural populations may vary in levels of species diversity, for instance after they experience disturbances that remove significant amounts of their genetic and species variation, including rare but functionally unique guilds. We investigated the influence of diversity on associated community function by propagating natural microbial communities from a traditionally fermented milk beverage diluted to various levels. Specifically, we assessed the influence of less abundant microbial types such as yeast and rarer bacterial types on community functionality and species sorting trajectories over approximately 100 generations. We observed repeatable changes in bacterial community compositions, metabolic profiles, and acidity related to starting diversity levels. The influence of a single ecological guild, yeast in our study, played a dramatic role on function, but interestingly not on long-term species sorting trajectories of the remaining bacterial community. Our results together evidence ecological selection on the microbial communities in our system and suggest an unexpected niche division between yeast and bacterial communities.

## INTRODUCTION

What factors affect dynamics of diversity in natural communities and how this links to community function has long been a central question in ecology (Cardinale et al., 2012; Gonzalez et al., 2020). Defining community function and community diversity at the level of species can take many forms. Community function has been estimated by quantifying parameters such as community productivity, overall metabolic output, stability or resistance against invasion (Aubree et al., 2020). Community diversity captures the living organisms present and their relative abundances. This diversity can be expressed at several levels of taxonomic richness; at the broader level of genus or species, or more narrowly at intraspecific diversity (species types). A general positive association between higher diversity and increased measures of community function has been shown (Balvanera et al., 2006), and it is typically interpreted to result from functional complementarity (Cardinale et al., 2007; Tilman, 1999) of collections of species with different niches (Cardinale et al., 2012).

Ecological processes such as species sorting – the sorting of variation at the level of species along an ecological or evolutionary timescale – may alter community diversity. For instance, when a particular species or type is better adapted to the new selective condition, they may increase in relative abundance (Vellend, 2010). The response to selection may depend on initial species diversity — both in the number of species present and their relative abundance. Species sorting may also affect overall community functioning, when species or types that increase or decrease in abundance are key to overall community function. While bodies of theory exist on the theme of species sorting in communities (Loeuille and Leibold, 2008), experimental tests are relatively few (Langenheder and Székely, 2011). Since most experimental studies on the influence of community diversity on community function focus on short-term ecological timescales of one or few generations (Aubree et al., 2020; Gonzalez et al., 2020), relatively little is known on how this influence changes over multiple generations (Fiegna et al., 2015).

As an experimental model system, the microbial communities of fermented foods provide a powerful method to study effects of selection on species diversity and on community function (Alekseeva et al., 2021; Wolfe, 2018; Wolfe and Dutton, 2015). In these communities, function is often quantified through changes in pH and through metabolic output measured as volatile compound production, contributing to aroma and taste, which can be further connected to known biochemical pathways (De Filippis et al., 2016; Krömer et al., 2004; Smid et al., 2005). Species diversity can be traced to the level of microbial groups known to be responsible for fermentation, such as lactic acid bacteria, acetic acid bacteria, and alcohol producing yeast. Metabolic functional groups can be considered equivalent to the classic ecological classification of species into guilds, lifeforms, or strategies (Louca et al., 2018). The well-studied metabolic, or fermentative, microbial groups of fermented foods are therefore relevant communities for studying fundamental questions in ecology.

Specifically, a traditionally fermented milk beverage from Zambia, Mabisi (Moonga et al., 2020; Schoustra et al., 2013), provides a model system to study the relationship between community diversity and function, and how this may change over repeated cycles of sequential propagation. The moderate diversity of Mabisi, composed of six to ten dominant lactic and acetic acid bacterial species, plus numerous other low abundance types (other bacterial species, yeasts, and viruses), further facilitates experimental design and analysis (Moonga et al., 2020; Schoustra et al., 2013). The defined and measurable functional properties of Mabisi, including metabolite profiles and acidity, frame investigations of the influence between diversity and community function. Four fermentative types are naturally present at various levels of relative abundance. These include: 1. alcohol producers, 2. alcohol consumers producing acetic acid, 3. homofermentative lactic acid producers (only lactic acid produced), and 4. heterofermentative lactic acid producers (lactic acid, ethanol, acetic acid, and carbon dioxide produced) (Gänzle, 2015). The presence and ratios of these four fermentative types, or ecological guilds, is expected to affect community metabolic profiles due to the distinctive metabolic capabilities each community member possesses.

Here, we present an experimental test of predictions on community functioning and species sorting using microbial communities at a range of diversity levels, which were repeatedly propagated for 16 cycles (approximately 100 generations) in milk in a laboratory environment. To evaluate the effect of diversity, we first serially diluted Mabisi microbial communities to generate four levels of initial diversity (Fig. 1). At propagation cycles 1, 5, and 17 we then measured metabolic output as a proxy for community function, and bacterial diversity by full 16S amplicon sequencing. At every second transfer, pH was also measured as indication of community function. With this we aimed to answer three main research questions: 1. Does altering initial diversity, with the associated loss of low abundance microbial types, influence community function? 2. And if so, are such shifts in community function stable over repeated propagation? 3. Does initial diversity influence bacterial species sorting trajectories over repeated cycles of propagation?

**Figure 1:**
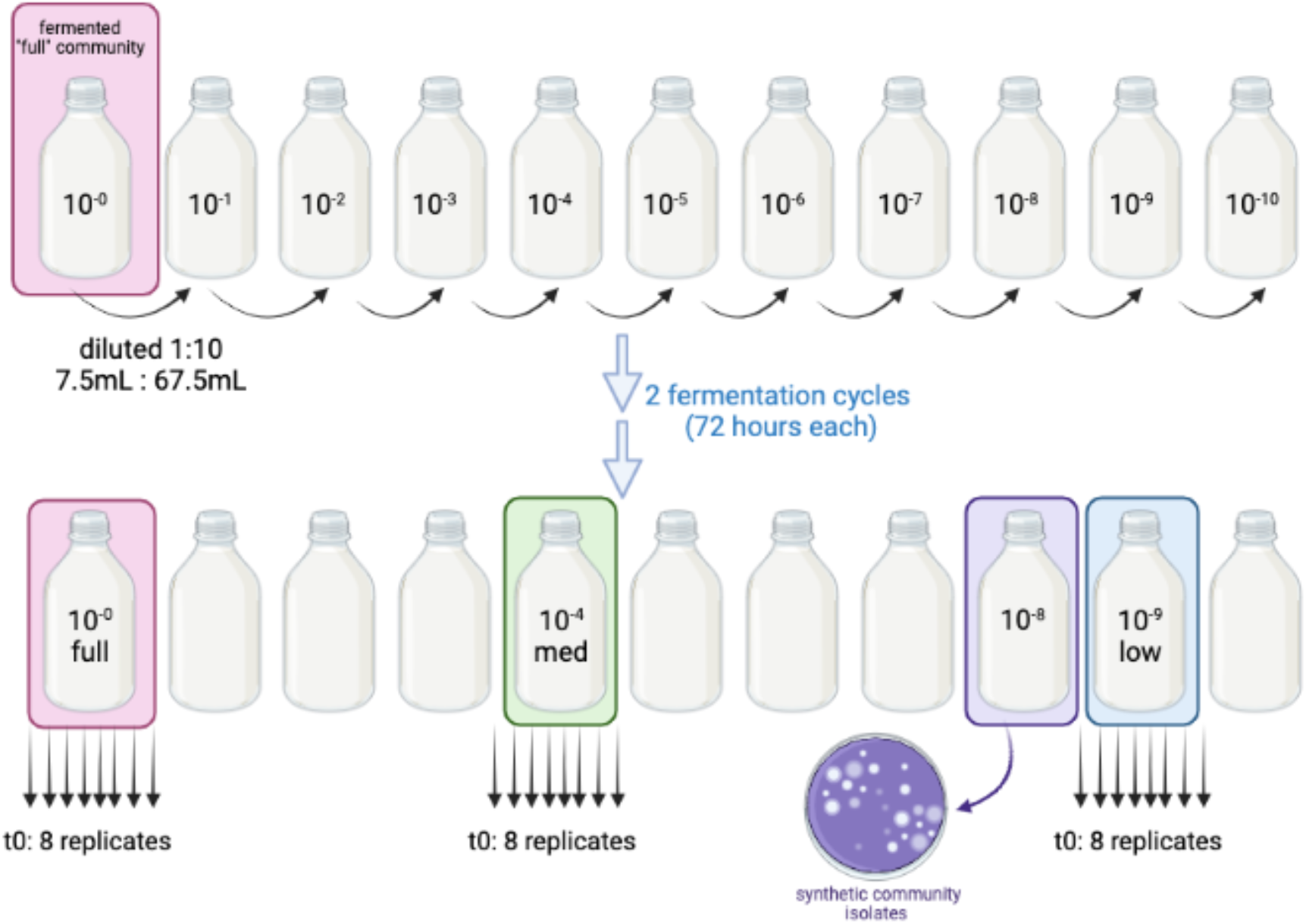
Experimental design. Dilution approach to create initial communities (t0), progressively removing rare types, arriving at four levels of initial species diversity (full, medium, and low diversity, and one synthetic community). From each diversity level eight replicates were then propagated for 16 transfers of 1% to sterile milk. Image created in BioRender.

## RESULTS

### Community function

Metabolic profiles of replicate communities at four levels of initial species diversity show that two main groupings of metabolic profiles persist across the three time points analysed, with full and medium diversity communities clustering together, versus low and synthetic (Fig. 2). The divide exists along principal component 1 (PC1) but not the second component (PC2). PC1 explains between 53.1% (transfer 5) to 65.5% (transfer 1) of the variation, while PC2 explains between 11.9% (transfer 1) to 19.6% (transfer 17). A division in metabolic profiles between full and medium communities is observed along PC2 at transfer 1 (Fig. 2a), disappearing at transfer 5 (Fig. 2b). The metabolic profiles of communities at the four diversity levels inconsistently exhibit relatively larger or smaller spread in data points across the time points but replicates of the synthetic community (purple ellipses) arguably show closer clustering in metabolic profiles throughout the experiment.

**Figure 2:**
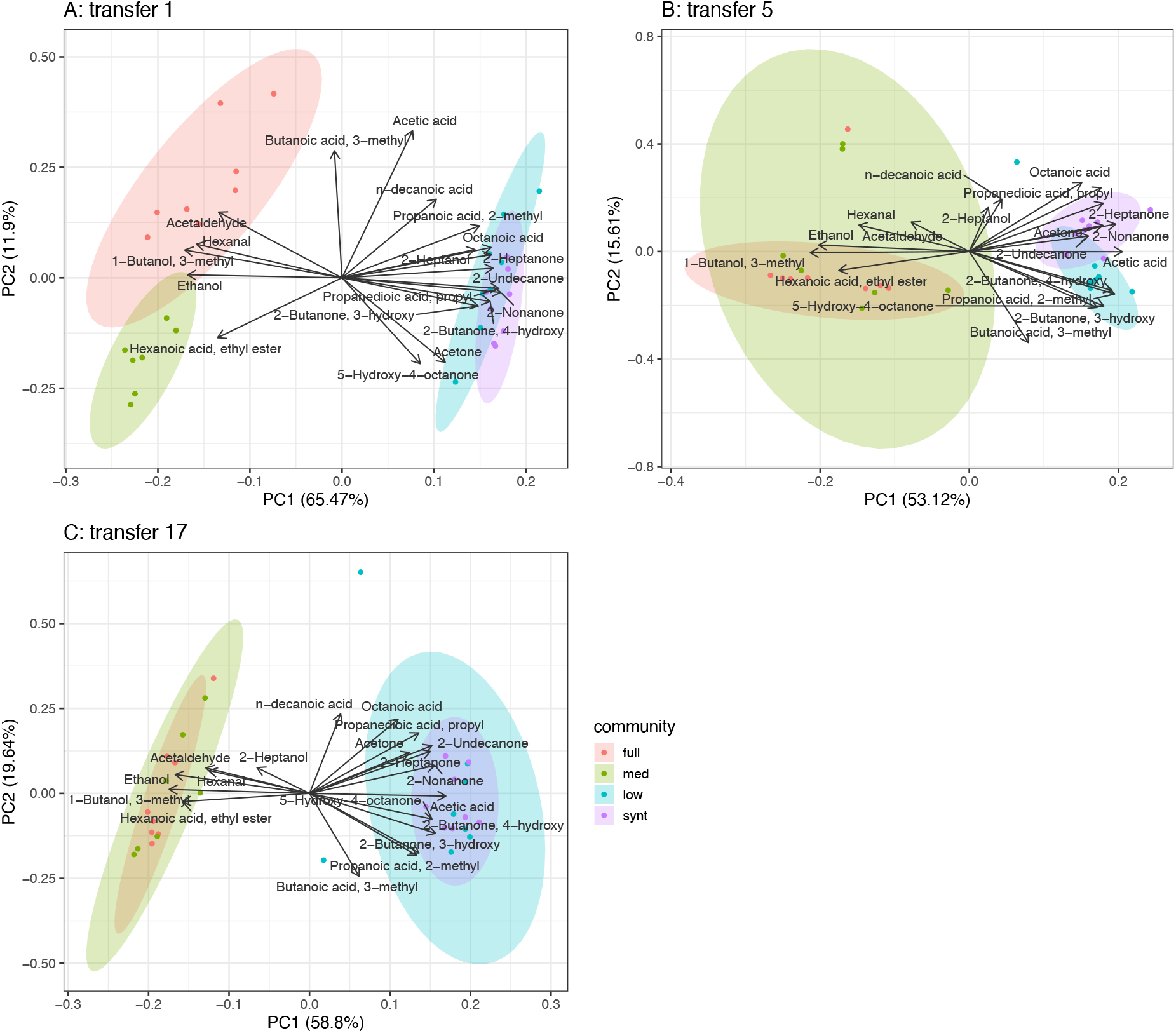
Diversity of metabolic profiles persist through propagation. Compounds associated with yeast metabolism primarily impact divisions in metabolic profile. PCA analysis of metabolic profiles compiled from 19 compounds using GC-MS analyses at transfer 1 (A), 5 (B), and 17 (C). Dots represent individual replicates. Transfer refers to the end product used for propagation that day (i.e., transfer 5 is end product from transfer 4). Ellipses are 95% percentile estimations. Direction of vectors indicate the compound’s contribution to either principal component, whereas the vector length indicates the amount of variation explained by the two plotted principal components. Yeast associated compounds: ethanol, acetaldehyde, hexanoic acid-ethyl ester, 1-butanol 3-methyl.

Figure 3 shows pH of cultures after a full cycle of growth over the course of the experiment. The overall pH differences across diversity treatments ranged between approximately pH 3.4 to 3.9. Except for transfer 1, low diversity and synthetic communities consistently display higher pH than full diversity and medium diversity communities. Low diversity communities maintained a higher pH until a drop between transfer 15 and 17 where they converge to a comparable value as the full and medium communities. Full diversity communities and medium diversity communities maintained similar pH throughout propagation. The pH continues to drop across all diversity treatments after transfer 5, most noticeably for the low diversity communities, while pH in synthetic communities remains significantly higher at transfer 17 (ANOVA: transfer 17 pH values, synthetic versus full community pH estimate = +0.151, p = 1.08e-07, F = 24.3 on 3 and 27 DF).

**Figure 3:**
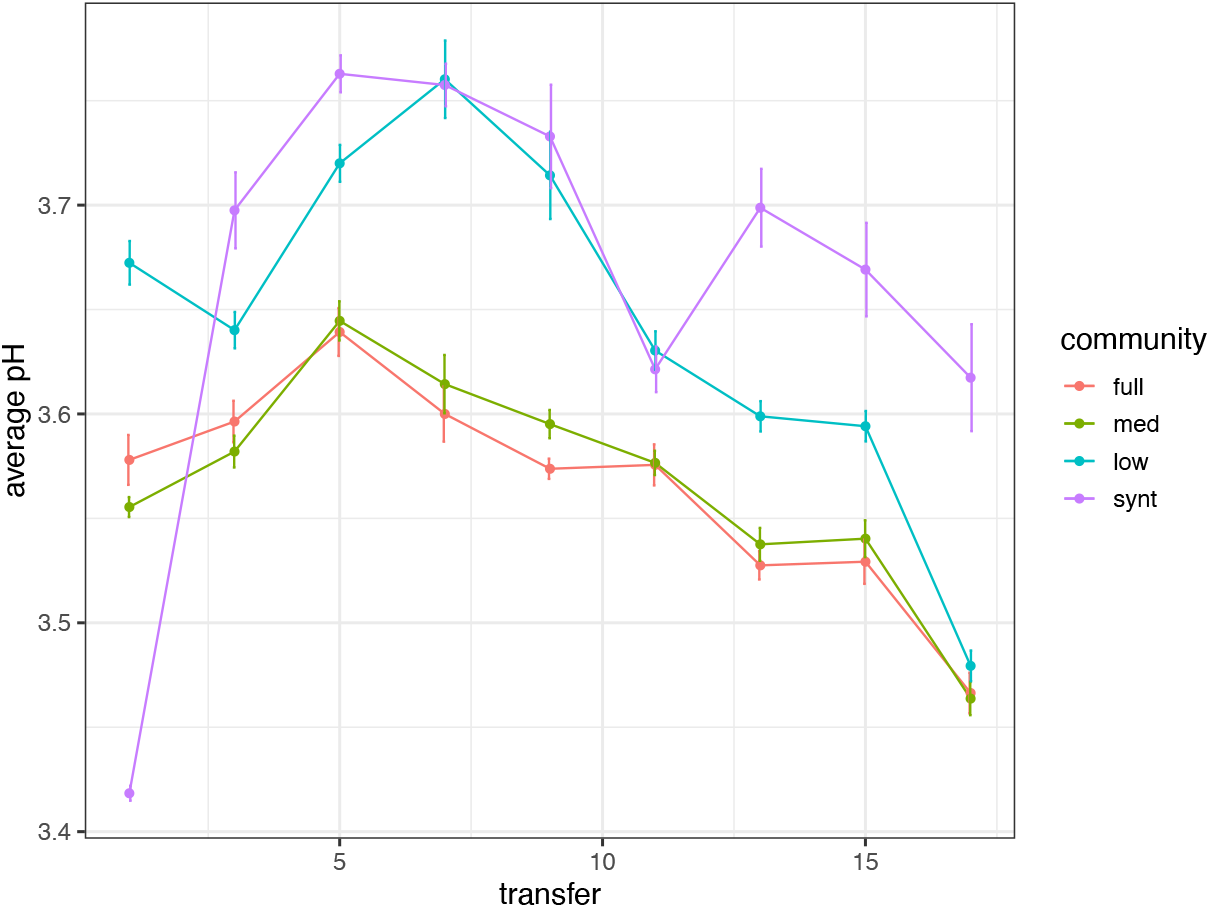
Loss of diversity initially increases pH, which continued propagation repeatedly decreases. Measures of pH of full diversity, medium diversity, low diversity and synthetic communities over 16 rounds of serial propagation. Average pH across replicates of the same community diversity level measured at every second transfer. Points show average value for the 8 replicate communities (except medium diversity treatment with 7 replicates), vertical bars show standard error of the mean.

### Bacterial species composition

Figure 4 shows relative abundance of bacterial species in each of the communities at transfer 1, 5, and 17. The results obtained for transfer 1 show the effect of serial dilution on bacterial community composition: compared to the full diversity communities, *Lactobacillus A* type (light turquoise) has a higher relative abundance in the medium diversity communities and is the dominant type in low diversity communities. This increase in relative abundance seen at transfer 1 of *Lactobacillus A* type diminishes over time, with *Lactobacillus B* type (medium blue) dominating in all replicates and initial diversity levels by transfer 17. However, the abundance of *Lactobacillus A* cluster remains comparatively slightly higher in low diversity communities at the end of propagation. This *Lactobacillus A* cluster was not included in the synthetic community but surprisingly appears in one synthetic community replicate at transfer 17, likely due to cross contamination during the serial propagation.

**Figure 4:**
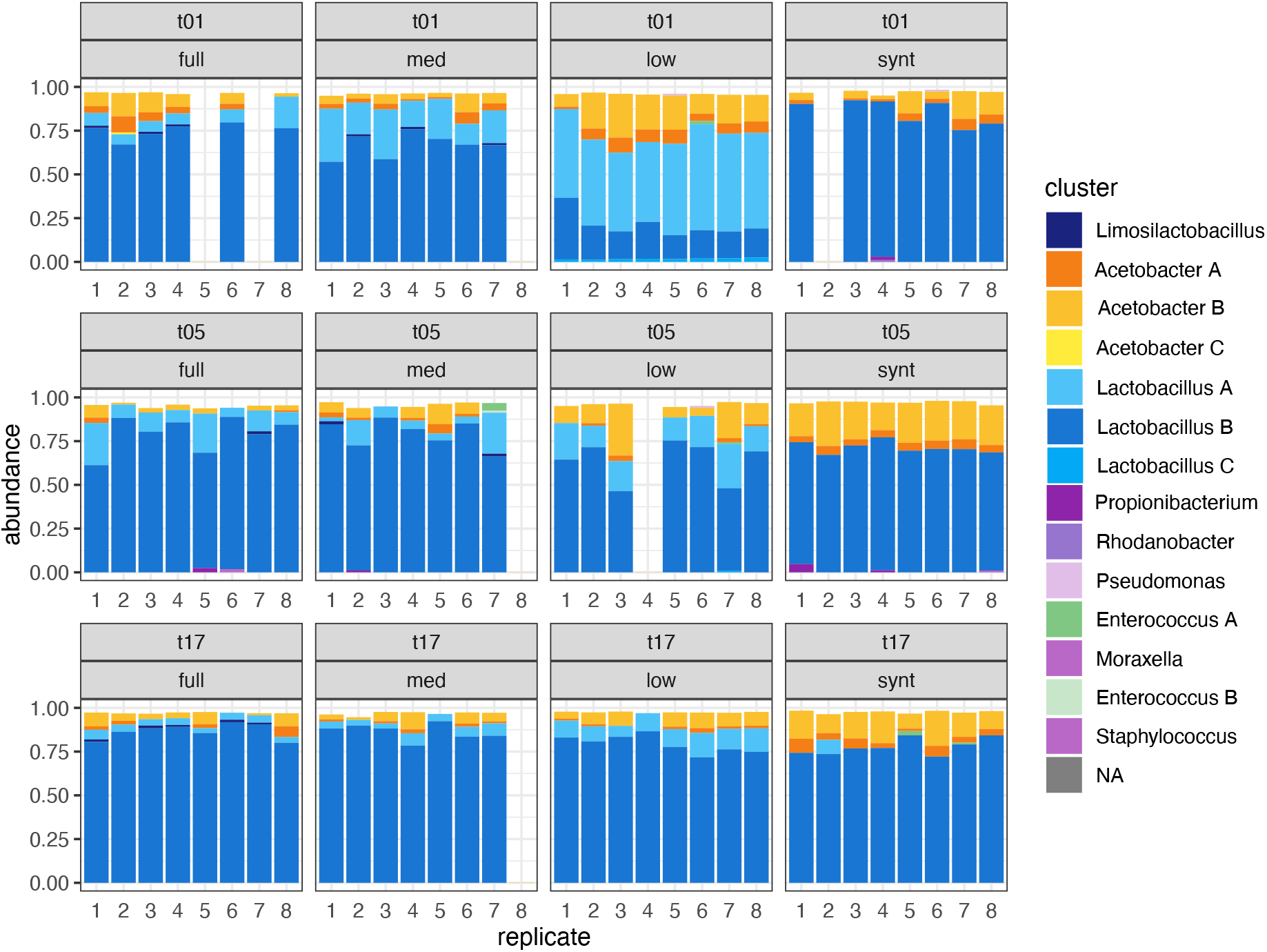
Starting diversity transiently alters bacterial community composition. Bacterial community compositions at transfers 1, 5, and 17. *Lactobacillus* clusters are shown in blue colour shades, *Acetobacter* in yellow-orange, *Limosilactobacillus* in dark navy blue, and low abundance contaminant types in green or purple. Clusters with <1% abundance were not identified nor plotted. Some replicate populations are missing due to insufficient DNA concentrations, resulting either from poor DNA extraction or library preparation. Medium community, replicate 8 was lost early in propagation and removed from all analyses.

Bacterial community composition significantly differed per diversity level and time point (PERMANOVA on community compositions, Bray method used: effect of initial diversity (p = 0.001, F = 70.9, DF = 3), transfer (p = 0.001, F = 55.1, DF = 2), diversity x transfer interaction (p = 0.001, F = 27.8, DF = 6)). Derived from the same data presented in Figure 4, NMDS plots of bacterial community compositions for transfer 1, 5, and 17 per initial diversity level (Fig. 5) demonstrate convergence across full, medium, and low diversity communities over time. Furthermore, final relative ratios of *Acetobacter A, Acetobacter B, Lactobacillus A, and Lactobacillus B* are very similar across all communities at transfer 17, apart from synthetic diversity ones (Fig. S3).

**Figure 5:**
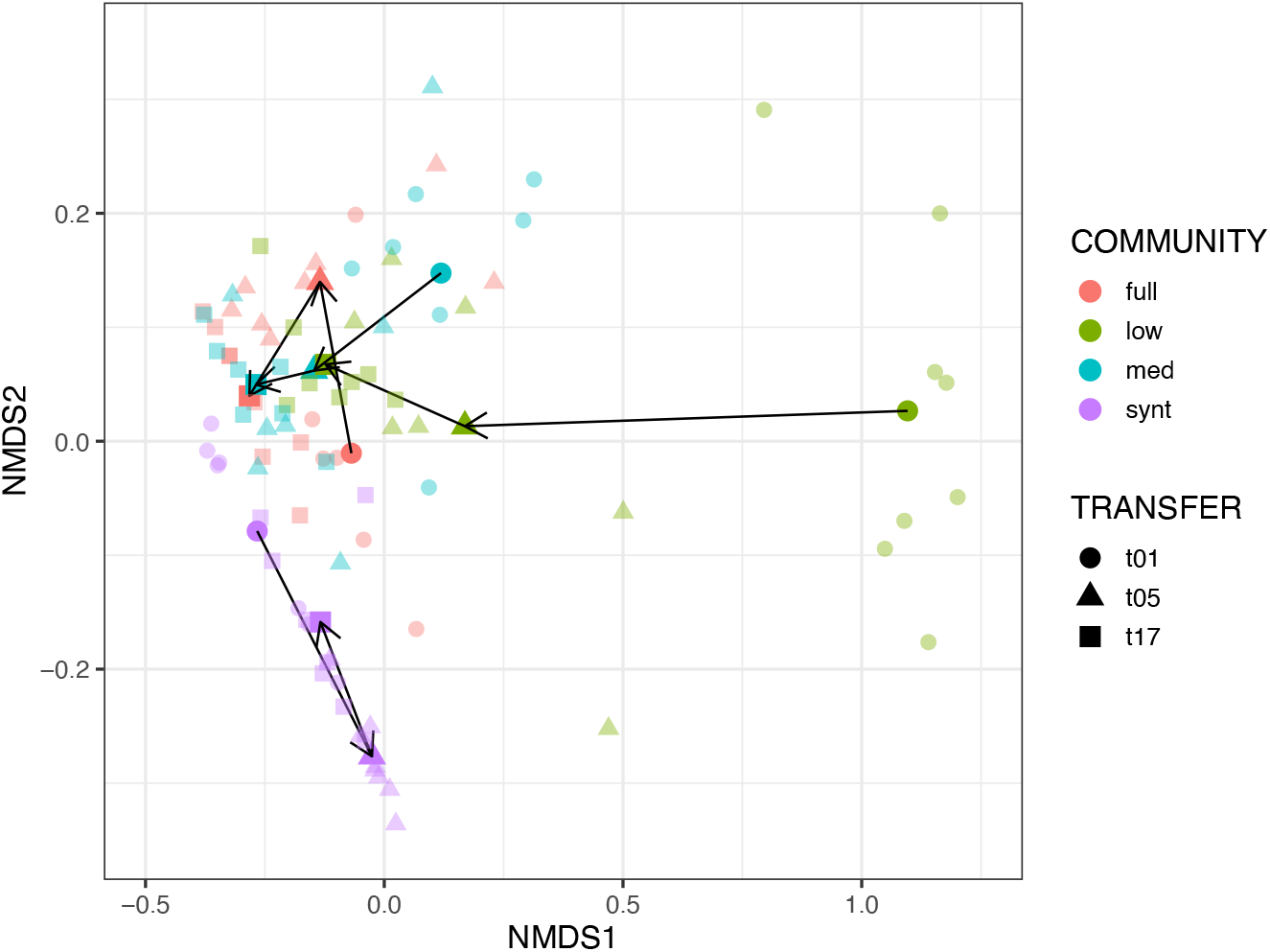
Dilution of diversity largely only transiently alters bacterial community composition. NMDS depiction of bacterial community compositions of diversity treatments at transfer 1, 5, and 17. Solid coloured points are centroids of data points for community*transfer grouping. Arrows connect centroids over time.

Variation in ratios of *Acetobacter* versus *Lactobacillus types* (yellow-orange versus blue colours, respectively) exist both between diversity treatment, as well as across replicates within a diversity treatment. *Acetobacters* are at highest relative abundance in the low diversity communities early in propagation, but synthetic communities demonstrate the highest proportions of *Acetobacter* types by transfer 5 and transfer 17. However, the relative abundances of *Acetobacter* versus *Lactobacillus* types do converge across full, medium, and low communities by transfer 17 (Fig. S2).

## DISCUSSION

In this study we assessed how progressively diluting a natural microbial community altered community function and how this function, as well as bacterial diversity, would subsequently change due to selection upon repeated cycles of propagation. By exploring the relationship between community function and diversity overtime, we observed repeatable changes of replicate lineages for metabolic profiles and acidity related to starting diversity levels of community compositions. These results show a clear division in metabolic profiles with full diversity and medium diversity communities on one hand, and low diversity and synthetic communities together on the other hand. This division was sustained throughout repeated rounds of propagation. Further, we found that changes in bacterial community composition (i.e., bacterial species sorting trajectories) over repeated cycles of propagation generally resulted in a convergence of bacterial communities to the same composition, irrespective of initial diversity level.

Our design was inspired by a specific prediction on community function at various levels of species diversity. Upon diluting the full community (10^0^) to medium (10^-4^) and low (10^-9^) diversity levels, then by using a synthetic community of five isolates from the community (Fig. 1), we progressively eliminated rare types (Table 1). Specifically, yeast was eliminated in low and synthetic communities (as visualised by eye and previous abundance estimates (Moonga et al., 2019; Schoustra et al., 2013)). The absence of yeast in the low and synthetic communities was predicted to have the strongest influence on metabolic profiles due to its divergent metabolic capacities versus background bacterial communities. High functional redundancy is observed in the bacterial part of the microbial community (Gralka et al., 2020; Wagg et al., 2019), thus we predicted that general acidification properties of lactic acid and acetic acid bacteria would be maintained in medium and low communities, regardless of the loss of yeast or rare bacterial types (Wagg et al., 2019; White et al., 2020). Regarding bacterial diversity, initial responses as well as the responses over multiple rounds of propagation to the loss of yeast could follow alternate outcomes, which in part may depend on the initial diversity level. On the one hand, niche space is freed upon community dilution and due to functional redundancy, the remaining bacterial community may partially restore metabolic capacities originally contributed by yeast. On the other hand, the niche space filled by yeast or other rarer types may be sufficiently unique and does not overlap with that of the remaining bacterial community.

**Table 1:** Presence/absence of fermentative types between diversity level treatments.

| Microbial type | Fermentative type | Diversity level |  |  |  |
| --- | --- | --- | --- | --- | --- |
|  |  | high | medium | low | synthetic |
| *Geotrichum* (yeast) | alcohol producer | + | + | - | - |
| *Lactobacillus* | homo fermentative | + | + | + | + |
| *Limosilactobacillus* | hetero fermentative | + | + | - | - |
| *Acetobacter* | alcohol to acetic acid | + | + | + | + |
| others & genotypes |  | + | +/- | +/- | - |

Although at minority abundances in typical Mabisi communities (Schoustra et al., 2013), yeasts contribute a unique metabolism since they are the sole alcohol fermentative type and are known to produce other distinct compounds such as methyl-esters. We thus predicted that presence of yeast in the full and medium diversity communities or, conversely, its absence in the low diversity and synthetic communities, would significantly influence community function. Initial metabolic profiles measured at transfer 1 provide support for this expectation, where alcohol and ester compounds typically linked to yeast metabolism are found. The presence of yeast in only the full and medium diversity communities is reflected by presence of ethanol, acetaldehyde, and hexanoic acid-ethyl ester among the main metabolites detected. Acetaldehydes and esters are a known products of yeast metabolism (Dzialo et al., 2017; Liu and Pilone, 2000). An additional compound with vector contributions towards the grouping of full and medium communities is 1-butanol-3-methyl, which is the breakdown product of leucine via the Ehrlich pathway found in various yeast species (Szudera-Kończal et al., 2020).

Heterofermentative types are also absent in low diversity and synthetic communities, whose presence is expected to alter metabolite profile; however, since this overlaps with the presence/absence of yeast in the communities, we are unable to disentangle which metabolite changes are specific to the presence or absence of heterofermentative types. Metabolic profiles are similar between full diversity and medium diversity communities, yet their bacterial compositions differ with losses of rare bacterial genotypes in medium communities, suggesting that these rare types do not significantly affect overall metabolite profiles. No effect, or a very weak effect, of rare genotypes on community function is further supported by the observation that low diversity and synthetic communities also show similar metabolic profiles despite different community compositions. Hence, in Mabisi communities, intraspecific bacterial genotype compositions do not appear to strongly influence metabolic profiles, suggesting functional redundancies in bacterial metabolic capacities. Overall bacterial community composition and function were similar regardless of loss of rare bacterial genotypes (i.e., full versus medium diversity, and low versus synthetic diversity). All diversity levels contained the two dominant fermentative types – homofermentative lactic acid bacteria (*Lactobacillus* A, B) and acetic acid bacteria (*Acetobacter* A, B) – which our data suggest are the core bacterial members for community function at the level of metabolite production, in addition to yeast. While yeast appear as a core microbial community member in our study’s Mabisi sample, interestingly, not all natural Mabisi products contain yeast (Moonga et al., 2020; Schoustra et al., 2013). Microbial community profiles of Mabisi samples vary across regions and processors in Zambia. There is past and ongoing research to link microbial community compositions to processing methods (Moonga et al., 2020) and consumer preferences.

At the level of acid production there exists great functional redundancy within the bacterial community, thus we predicted that pH would not differ greatly between the initial diversity levels. We found support for this prediction; however, a very slightly higher pH was observed for the synthetic, hence lowest diversity, community. This modest increase suggests that the presence of yeast and rare bacterial genotypes does not impact community acidification. A dynamic jump in pH in synthetic diversity communities between transfers 1 and 3 is not surprising since this community was more naïve in its member structure and substrate (these isolates had been pre-cultured in MRS broth). The range in pH between diversity levels is modest (~pH 3.4 to 3.85) but may still represent a selective force on the microbial communities. Mabisi acidity increases quickly during fermentation, with the most notable pH drop between 24 to 48 hours (Groenenboom et al., 2020). Our observed trend of continually decreasing pH may be explained by the preferential propagation of, and thus selection for, persisting abundant types in the acidic conditions created after 72 hours of fermentation. Increased acidity is a common outcome of bacterial community domestication and repeated back-slop propagation of fermentation starter cultures (Bachmann et al., 2011; Spuś, 2016; Van Kerrebroeck et al., 2016). It would be interesting to test how much lower the pH would reduce with further propagation and when or if pH stabilisation would be reached. Evolving communities could be improving resource use, consequently excreting more metabolites due to larger supported population sizes, further lowering the pH over additional repeated transfers.

Species profiles of the bacterial communities at transfer 1 show a clear signature of the progressive serial dilution of diversity into full diversity, medium diversity, low diversity and synthetic communities. Although progressive removal of microbial types across our starting communities affected initial and final metabolic function, it interestingly did not have a large effect on the outcome of species sorting trajectories of the remaining bacterial community over 16 cycles of propagation. This is especially true when focusing on the relative ratios of the major bacterial guilds - acetic acid versus homofermentative lactic acid bacteria, which were surprisingly uniform across full diversity, medium diversity, and low diversity communities after 16 cycles of propagation. If replicate communities of higher diversity diverged more so in their compositions overtime, then our results would have aligned with the theory that “diversity begets diversity” via greater niche construction (Madi et al., 2020). Reversely, our results could have exhibited replicate communities of lower diversity diverging from one another, which would support the concept that greater diversity restricts possible trajectories due to less available niche space (van Moorsel et al., 2021). The minimal effect of diversity on sorting trajectories in our experiment could be explained by high functional redundancies in the major bacterial types (i.e., acetic acid and lactic acid bacteria).

Most surprising was the seeming lack of influence of yeast on final bacterial compositions; all diversity levels were dominated by *Lactobacillus* B and had similar ratios of *Acetobacters* and *Lactobacilli*. The convergence of bacterial community compositions, regardless of the presence of yeast communities, suggests yeast and bacteria exist in sufficiently unique niches of resource use. Evidence for strong division in resources and hence lack of influence of yeast on bacterial community composition in a fermented food was surprising and contrasts against previous findings showing metabolic associations between the two (Blasche et al., 2021; Mendes et al., 2013; Ponomarova et al., 2017; Suharja et al., 2014; Xu et al., 2021). However, these investigations mostly focus on *S. cerevisiae* and specific *Lactobacilli* species or strains; the yeast in our system is identified as *Geotrichum candidum*. If *Lactobacilli* versus *Acetobacters* had differing metabolic associations with yeast, we would expect shifts in their relative abundances following the removal of *G. candidum*, but this was not observed. We can thus hypothesise that in the Mabisi system, yeast do not affect bacterial growth by consuming end products of bacterial metabolism such as lactate, thus are simply secondary in the metabolic route. In addition to more cross-feeding like interactions, yeast and bacteria could be in resource competition, in which case the removal of yeast would open niche space for certain bacterial types. Bacterial types whose niche space previously overlapped more with yeast, or were poor competitors, would expectedly establish at higher abundances in the absence of yeast; this is not what we observe. Overall, our results support that amongst bacterial types in our community, they are all in similar, at most very weak, resource competition with yeast since bacterial community compositions converge across all diversity levels regardless of yeast’s presence. A next step to further elucidate the influence of yeast on community function in Mabisi is to add isolated *G. candidum* to the low and synthetic communities, then compare metabolic profiles and acidity. The reverse direction could also be taken, by eliminating yeasts from the full and medium communities with fungicide, which would maintain rare bacterial types.

We interpret the observed repeatability between replicate lineages and convergence in bacterial communities to be evidence of ecological selection (Vellend, 2016, 2010) in our study system. In evolutionary biology research, repeated or convergent ratios of genotypes under a common environment is interpreted as evidence for adaptive selection (Hughes, 1999); we take the same interpretation for our results but at the ecological level of ratios of identified bacterial types. However, we currently can only hypothesise about the sources of ecological selection in our study, with possibilities including our chosen laboratory conditions (temperature, fermentation time and associated acidity, supermarket milk content) and unspecified biotic species interactions. While prokaryotes and bacteriophage are known members of the microbial community of our natural experimental system (Schoustra et al., 2013), we have focussed our analyses on species sorting in the bacterial part of the microbial community since bacteria are present in all communities regardless of starting diversity (i.e., dilution) treatment.

Studying functionality responses on longer time scales after an erosion of species and/or genotypic diversity remains an underexplored topic that we made initial steps here to explore. The ecological processes of species sorting in a community can be paralleled to evolutionary dynamics of genotypic changes within a species (Vellend, 2016, 2010). Understanding how changes across hierarchical levels — from genes, to communities, to ecosystems — influence one another is undoubtedly complicated and it merits further steps to elucidate. In this study we focused on ecological processes without investigating genotypic changes. However, we foresee exciting future avenues of research to unravel the role of ecology versus evolution in altering communities and their functionality. Creating a synthetic community comprised of four fermentative types that grow in a defined media, combined with whole genome sequencing and/or metagenomics could allow identification of novel genotypes, and is an exciting avenue to better explore ecological-evolutionary dynamics.

## METHODS

### Preparing communities (t0)

A fermented Mabisi sample containing its full microbial community was serially diluted up to a factor of 10^-10^ using 7.5 mL culture with 67.5 mL UHT full fat milk (Jumbo brand, Houdbare Volle Melk). Only one replicate dilution series was performed, creating one culture bottle for each dilution level. The entire volumes of these dilutions were then fermented for 72 hours at 28 °C, then 0.75 mL of final culture was transferred to 75 mL of fresh milk for a second 72-hour fermentation at 28 °C. Cultures were swirled well to mix, especially at the air-liquid phase, before transferring. After the second fermentation, the Mabisi products created from each serially diluted community were compared. A thickened, fermented product was observed up to the 10^-9^ diluted community and yeast were visibly present at the air interface up until the 10^-4^ diluted community. Visual observations informed the choice of three levels of diversity to use in the main evolution experiment: full community (10^0^ dilution), medium (10^-4^ dilution), and low (10^-9^ dilution). Initial T_0_ communities were archived from the final product after two fermentation cycles without glycerol at −20 °C and with glycerol at −80 °C (1 mL culture + 0.5 mL 85% glycerol). Yeast and whey production was therefore seen only in medium and full communities.

A synthetic community of five individual isolates was also created as a fourth diversity level (Table S1). A Mabisi community that had been diluted to 10^-8^ and undergone two 72 hours fermentation cycles at 28 °C was diluted and grown aerobically at 28 °C on MRS (de Man, Rogosa, Sharpe) agar. Initially eight unique appearing morphotypes (labelled A-H) were selected from the agar plate for investigation. The community was also plated on PCA (plate count agar) and M17 agars, but the MRS provided the clearest and most diverse collection of colony types. Colonies were grown for five days in MRS broth and archived at −80 °C (1 mL culture + 0.5 mL 85% glycerol). The eight colonies were assessed for their growth in MRS broth, morphotype on MRS agar, API metabolism test (BioMérieux), and ability to acidify milk in isolation. Using the mentioned assessments, a final five colonies (A, B, C, F, G) were chosen based upon being the most dissimilar (confident at least three were unique). Each colony was streaked twice on MRS agar and inoculated five days growth at 28 °C in MRS broth (1.5 mL broth in 24 well plate). The cultures’ optical density was measured (OD_600_) and then combined in volumes containing equal cell densities of each. This mixed culture formed the synthetic T_0_ diversity community and was archived at −80°C (1 mL culture + 0.5 mL 85% glycerol). Later Sanger sequencing of the full 16S gene (27F, 1492R primers) with Blast search in the NCBI database was unable to decipher the colonies to the species level (Table 1), as top identifications were all a >99% match. However, the 16S sequences indicated that at least three unique types were present – two *Acetobacter* and one *Lactobacillus*.

### Transferring and archiving

For each diversity level (full, medium, low, synthetic) eight replicate lines were inoculated using their respective T_0_ frozen cultures (1.5 mL total, with glycerol) in 75 mL of milk (total 32 evolution lines, plus one uninoculated milk as a negative control). Samples underwent repeated 72-hour fermentation cycles at 28 °C for 16 transfers (approximately 100 generations). For logistical purposes, after every second transfer final cultures were inoculated into fresh milk and stored at 4 °C for 24 hours before being moved to 28 °C. Therefore, a “true” cycle was completed after every second transfer (i.e., 7 days). Fresh samples of final products were archived at transfers 1, 2, 3, 5, 11, and 17, with the following archived: with glycerol at −80°C (1.27 mL culture + 0.63 mL 85% glycerol), and without glycerol at −20°C for DNA analysis and GC-MS analysis (~9 mL). Please note that archive labelling was done as follows: t5 = final product from t4 (i.e., the Mabisi culture used to inoculate at transfer 5), t17 = final product from t16. Due to an error in transferring, the M8 line (medium diversity, replicate 8) was lost early in the experiment, thus it is excluded from all analyses.

### GC-MS Analysis

Larger tubes of samples without glycerol at −20°C were defrosted at 4 °C, thoroughly mixed, 1.8 mL pipetted into headspace vials, then stored at −20 °C until analysis. A sample of 1.8 mL Jumbo Brand Volle Melk and 1.8 mL Jumbo Kefir Naturel were included with every time point as controls. After incubating for 20 minutes at 60 °C, a SPME fibre (Car/DVB/PDMS, Suppelco) extracted volatiles for 20 minutes at 60 °C. Volatiles were desorbed from the fibre under the following conditions: Stabilwax-DA-Crossbond-Carbowax-polyethylene-glycol column (2 min), PTV split mode at a ratio of 1:25 (heated to 250 °C), helium carrier gas at 1.2 mL/min, GC over temperature at 35 °C (2 min) raised to 240 °C (10 C/min), kept at 240 °C (5min). Mass spectral data was collected over a range of 33-250 m/z in full scan mode with 3.0030 scans/seconds. Results were analysed with Chromeleon 7.2 CDS Software (ThermoFisher) where the following signal peaks were identified as volatile metabolites according to their elution time and mass spectral data: acetaldehyde; acetone; ethanol; hexanal; 2-heptanone; 1-butanol, 3-methyl; hexanoic acid, ethyl ester; 2-butanone, 3-hydroxy; 2-heptanol; 5-hydroxy-4-octanone; 2-butanone, 4-hydroxy; 2-nonanone; acetic acid; propanoic acid, 2-methyl; 2-undecanone; butanoic acid, 3-methyl; propanedioic acid, propyl; octanoic acid; and n-decanoic acid. MS quantification peak counts were exported to Excel.

Data was first normalised by compound using the calculation 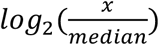, where x is the quantification peak count for a given compound in a given sample (i.e., area under compound peak), and the median ion count across all samples for that compound. Due to the viscous nature of Mabisi, especially the synthetic community samples at later time points, accuracy and reliability of volumes were questionable. For example, pipetting the highly viscous and thick synthetic diversity samples was very difficult, creating probable variation in sampling volumes. To be conservative, data was therefore further standardised by sample using the equation 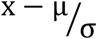, where μ is the mean quantification peak count and σ the standard deviation (following normalisation by compound) across compounds for a given sample. We realised that this removed all quantitative information about total aroma intensity between samples or diversity levels and left information only about relative compound peak heights. However, it was the justifiable approach considering the likely inaccuracies of sample volumes.

### DNA extraction

Adapted from Groenenboom et al. 2020 and Schoustra et al. 2013 (Groenenboom et al., 2020; Schoustra et al., 2013). A sample of 1.8 mL of fermented milk was spun down (2 minutes, 12000 RPM), then the supernatant and curd removed with a sterile scoopula. Cells were re-suspended in a mix of 64 μL EDTA (0.5 M, pH 8), 160 μL Nucleic Lysis Solution, 5 μL RNAse, 120 μL lysozyme [10 mg/mL] and 40 μL pronase E [10 mg/mL] and incubated for 60 minutes at 37 °C with agitation of 350 RPM. Cells were dislodged by manually flicking the tube occasionally during incubation. This was to improve mixing with suspension mixture. Bead beading was then performed for 3 minutes (1 minute, 5 minute rest, repeated three times) with sand sized beads, then 400 μL ice-cold ammonium acetate (5 M) added, and the mixture immediately cooled on ice for 15 minutes. The mixture was spun down (13000 rpm x g, 4 min) and 700 μL of supernatant transferred to a new 1.5 mL tube. Equal volume of phenol (700 uL) was added, the tube vortexed, and then spun down (6 minutes, 12000 RPM, 4 °C). 300 μL of supernatant was transferred to a new tube, 300 μL chloroform added, then was vortexed and its content spun down (2 minutes, 12000 RPM). From here, 300 μL of supernatant was transferred to another new tube, 400uL of 2-isopropanol added and vortexed. This mixture was left at −20°C overnight to precipitate.

Following overnight precipitation, the tube was spun down (13000 rpm, 4 °C, 15 minutes), then the supernatant carefully poured out so that the DNA pellet remained. 1mL of 70% cold ethanol was added to the tube and then spun down (10 min, 12000 rpm, 4 °C). The supernatant was carefully poured out and the DNA pellet washed again with 1mL 70% cold ethanol, spun down, and supernatant poured out. The DNA pellet was left to dry at 37 °C for 5 min, then dissolved in 20 μL of TE buffer (pH 8.0). A brief incubation (<30 min) at 37 °C improved dissolving the DNA pellet. Extracted DNA samples were stored at −20°C. Extracted DNA was measured on Qubit using dsDNA High Sensitivity kit, and a new diluted sample made with DNA concentration 0.5 ng/μL.

### Nanopore MinIon protocol

Adapted from Beekman et al. 2022 (Beekman et al., 2022).

DNA concentrations in these steps were always measured on Qubit 2.0 fluorometer using dsDNA High Sensitivity Assay kit (Thermo Fisher).

#### Determination of number of PCR cycles

An approximate appropriate number of PCR cycles was first determined on just three samples (1-F1, 1-L1, 1-S1), using 13, 16, 18, and 25 cycles. Primers, amounts of reagents, and PCR settings were as described below in “Tailed PCR reaction”. End products were visualised on 1% agarose gel and the minimal number of cycles decided based upon the fewest number of cycles where a sufficient band was seen (determined to be 23-30 cycles).

#### Tailed PCR reaction

Nanopore tailed forward: 5’ TTTCTGTTGGTGCTGATATTGC-[27F] 3’ Nanopore tailed reverse: 5’ ACTTGCCTGTCGCTCTATCTTC-[1492R] 3’ 27F: 5’ AGA GTT TGA TCC TGG CTC AG 3’ 1492R: 5’ TAC GGY TAC CTT GTT ACG ACT T 3’

The first step in Nanopore sequencing was a PCR reaction using Nanopore specific tailed primers. The specific number of cycles used for each sample is seen in Table S2. Two positive controls were included – the ZymoBIOMICS Microbial Community DNA Standard D6305, and a previously sequenced Mabisi sample (Groenenboom et al., 2020). The tailed primer PCR reaction was as follows:

**Tailed reaction reagents:**

1 uL – DNA [0.5ng/μL]
12.5 uL – Phusion High Fidelity PCR 2X master mix (ThermoFisher)
1.25 uL – forward tailed primer [10uM]
1.25 uL – reverse tailed primer [10uM]
9 uL – MilliQ water

**Tailed cycle conditions:**

98 °C 10 sec
98 °C 5 sec (~ 25X, see Table S2 for specifics)
57 °C 5 sec (~ 25X)
72 °C 30 sec (~ 25X)
72 °C 1 min
12 °C infinity

The tailed PCR reaction was performed another two times, resulting in three separate tailed primer PCR products per sample. Placement of PCR tubes in machine was adjusted for each PCR to avoid edge effects. Each amplified DNA sample was all visualised on 1% agarose gel to confirm successful amplification, then 8 μL of each PCR reaction were combined. A total of 24 μL of amplified DNA per samples was used for the clean-up.

#### PCR Clean-up

For each sample, the 24 μL of amplified DNA was cleaned with 24 μL of homemade SPRI beads (i.e., 1:1 ratio) (1 ml Sera-Mag SpeedBeads (Cytiva, Marlborough, MA, USA) cleaned and dissolved in 50 ml end volume containing 2.5 M NaCL, 20 mM PEG, 10mM Tris-HCL and 1 mM EDTA) and eluted into 20 μL of MilliQ water. DNA concentration of cleaned amplicons was measured using Qubit 2.0 Fluorometer. A new dilution of 15 μL of the cleaned, amplified 16S PCR product was made into a new diluted sample with DNA concentration 0.5 nM.

#### Barcoding

The PCR for each sample was barcoded to enable pooling using the PCR Barcoding Expansion 1-96 Kit (Oxford Nanopore Technologies). Reaction volumes were adapted from the Nanopore barcoding protocol to save in reagents used. The reaction was as follows with a unique barcode per sample:

**Barcoding PCR (per reaction/sample):**

0.3 μL barcode (Oxford Nanopore Technologies)
7.2 μL [0.5nM] cleaned PCR
7.5 μL LongAmp Taq 2x Master Mix (New England Biolabs)

**Cycle conditions:**

95 °C – 3 min (x1)
95 °C – 15 sec (x16)
62 °C – 15 sec (x16)
65 °C – 1.5 min (x16)
65 °C – 2 min (x1)
4 °C – infinity

Before pooling of the PCR barcoded products, each was visualised on 1% agarose gel. All samples were combined with 2 μL, apart from samples with fainter bands that were subjectively determined to require 3 μL (1-F1, 1-S1, 5-L1), 4 μL (5-S1, 1-M5), or 5 μL (1-S2, 1-F5, negative control). The pooled sample (total volume = 208 μL) was cleaned using homemade SPRI beads in 1:1 volumetric ratio and eluted in 200 μL of milliQ water. The DNA concentration of the cleaned, pooled sample was measured using Qubit fluorometer.

In 47 μL of milliQ water, 1 ug of the barcoded, pooled, cleaned library was prepared. From here, the library was repaired, end-prepped and adaptor ligated according to the Oxford Nanopore Technologies PCR barcoding (96) amplicons (SQK-LSK109) protocol, version “PBAC96_9069_v109_revO_14Aug2019”. Reagents used were NEBNext FFPE DNA Repair Buffer (E7181A), NEBNext FFPE DNA End Repair Mix (E7182A), NEBNext Ultra II End Prep Reaction Buffer (E7183A), and NEBNext Ultra II End Prep Enzyme Mix (E7184A) (New England Biolabs).

The prepared library was loaded on a SpotON Flow Cell (FLO-MIN106D) on a MinION MK111775 sequencing device (Oxford Nanopore Technologies). Basecalling was performed shortly after using Guppy software version 6.2.4+a11ce76.

Barcodes removed due to too few reads were bc16 (1-S2), bc43 (5-L4), bc49 (1-F5), bc73 (1-F7). Also removed from final analyses were bc90 (neg), bc94 (zymo), bc86 (positive control, DNA isolated from Groenenboom et al. 2022 (Groenenboom et al., 2022)). Barcodes with fewer reads, but still sufficient (~7000 reads) were bc8 (5-S1), 50 (1-M5), 61 (1-F6), 64 (1-S6).

### Bioinformatics

To assign 16S sequences to taxonomic identity, we first downloaded the SILVA reference database (Quast et al., 2013). As we were interested in variation above the species level, a custom database was produced using vsearch to cluster 16S sequences at a 95% similarity threshold (Rognes et al., 2016). Reads were aligned to this database using minimap2 (Li, 2018), and the best matching cluster was assigned as the taxonomic identity for each read. Details of sequence compositions for main clusters are found in Table S3 excel file. Clusters to which less than five reads matched were assumed to be trace contamination and discarded.

The overall frequency of clusters with <1% abundance is similar across diversity levels, as evidenced by the amount of “missing reads” to complete 1.00 abundance on bar plot (i.e., blank space at top of bar plot). Since full communities and synthetic communities have similar frequency of the rarest clusters, they are likely present due to general contamination and not true diversity being removed through our analyses. Unexpected “external” reads were identified (purple and green colouring) but are at reasonable levels as expected from low level laboratory contamination amplified by repeated PCRs during Nanopore sequencing.

We used the SILVA database reference sequences to identify types of bacteria. Our bioinformatic analyses were able to identify *Lactobacillus* and *Limosilactobacillus* types to the species level, with all clusters containing nearly 100% of sequences with same species identification (Table S3 excel file). This was not true for *Acetobacter* types, where clusters included sequences matching to several *Acetobacter* species and these species matched to multiple clusters. For example, sequences labelled as *A. lovaniensis* are included in *Acetobacter A, B*, and *C* clusters. Hence, for the purpose of our study we only classified into general genus level groups for *Lactobacillus, Limosilactobacillus, and Acetobacters*. Our bioinformatic analysis could classify *Lactobacillus* and *Limosilactobacillus* clusters as homo or heterofermentative, respectively. The very rare *Limosilactobacillus* type detected in only some full and medium communities, is the sole identified heterofermentative type -*Limosilactobacillus fermentum*. The yeast present in the Mabisi communities used in this study was isolated and identified as *Geotrichum candidum* using amplification and sequencing of the ITS (internal transcribed spacer) region (top match GenBank: MK967716.1). Yeast was visibly seen growing at the air interface only in the full diversity and medium diversity communities.

## AUTHOR CONTRIBUTIONS

Experimental design and conceptual ideas were developed by AML and SS. Experiments and data collection were performed by AML with help of Dr. Francisca Reyes Marquez. Data analysis was completed by AML and BA, with BA creating the bioinformatic analysis pipeline. Manuscript written by AML and SS with editing support of BA and EJS.

## ACKNOWLEDGMENTS

We greatly thank Dr. Francisca Reyes Marquez for her laboratory expertise and support with DNA extractions and Nanopore MinIon sequencing protocols. We also thank Judith Wolkers-Rooijackers for running the GC-MS machine and helping with its associated data analysis. Thank you as well to Mariska Beekman for sharing her experience with Nanopore sequencing and its bioinformatic analysis.

## DATA AVAILABILITY

Source data files and codes available on GitHub (amleale/diversity_function_mabisi): https://github.com/amleale/diversity_function_mabisi.git. Sequence data will be uploaded to NCBI upon acceptance:

## SUPPLEMENTARY

**Figure S1:**
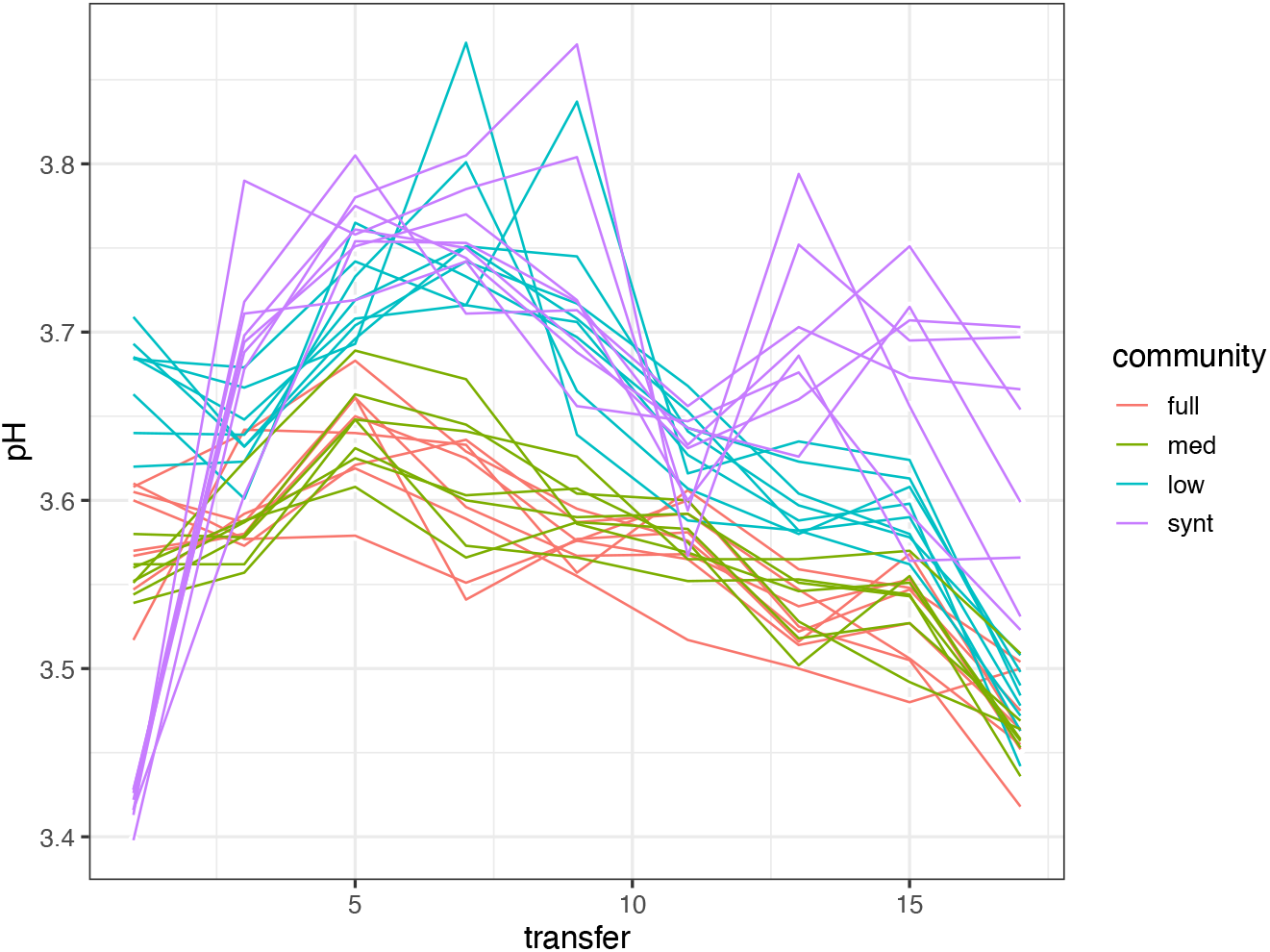
Measured pH at every second transfer for individual replicate communities over 16 rounds of propagation.

**Figure S2:**
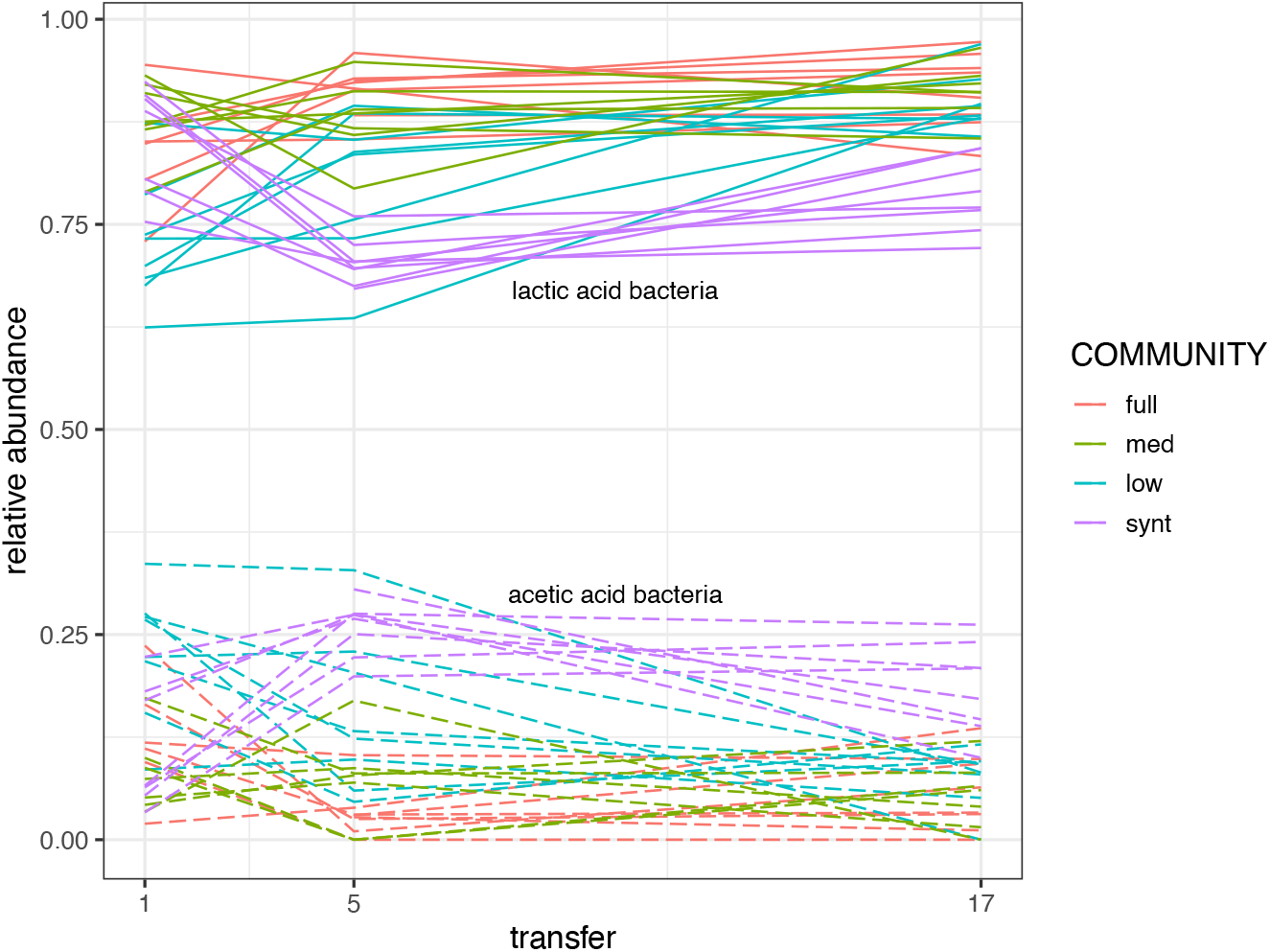
Community compositions become more similar over time, with the exception of synthetic community with more acetic acid bacteria. Relative abundance of lactic acid and acetic acid bacteria overtime. Lines follow each replicate over time. Solid lines: lactic acid bacteria, dashed lines: acetic acid bacteria. Lactic acid bacteria grouping: *Limosilactobacillus, Lactobacillus A, B, and C*. Acetic acid bacteria grouping: *Acetobacter A, B, and C*. Four replicate populations missing as in Figure 4.

**Figure S3:**
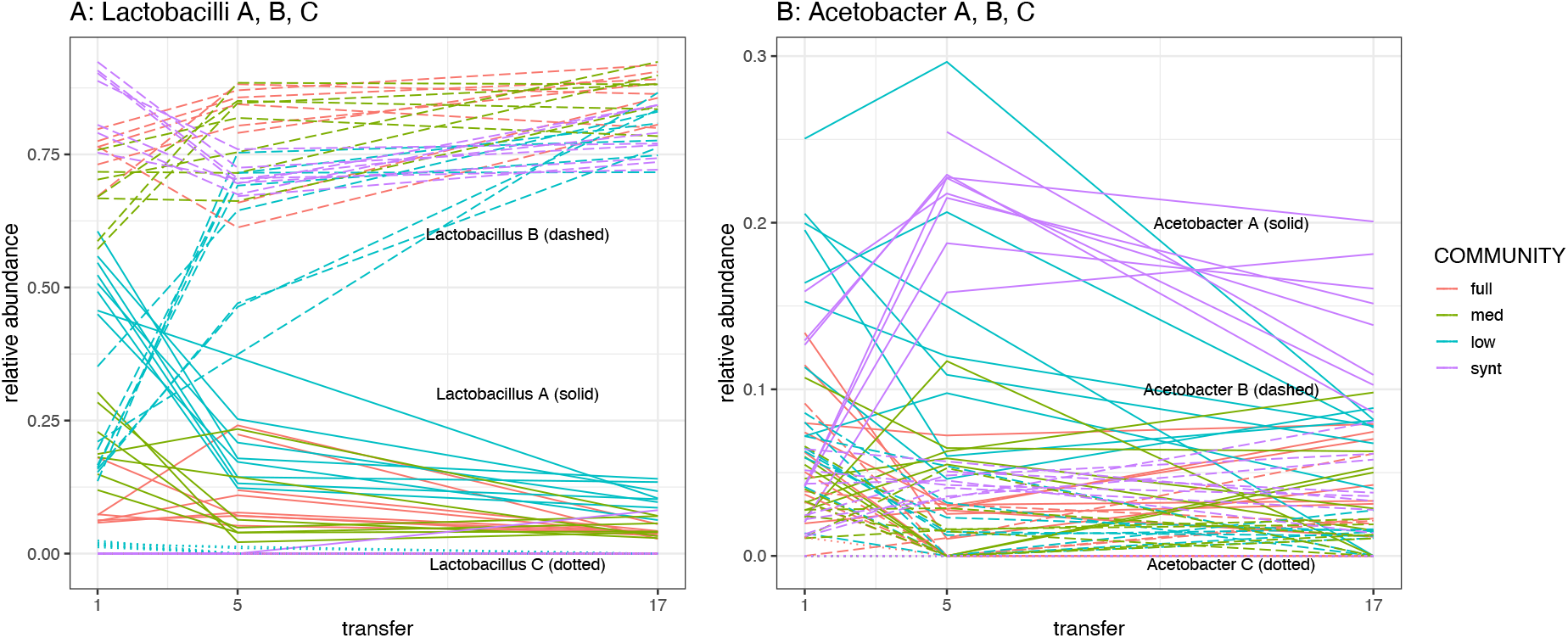
Relative abundances of lactic acid and acetic acid bacterial types converge overtime. Lines follow each replicate overtime. Four replicate populations missing as in Figure 4.

**Table S1:**
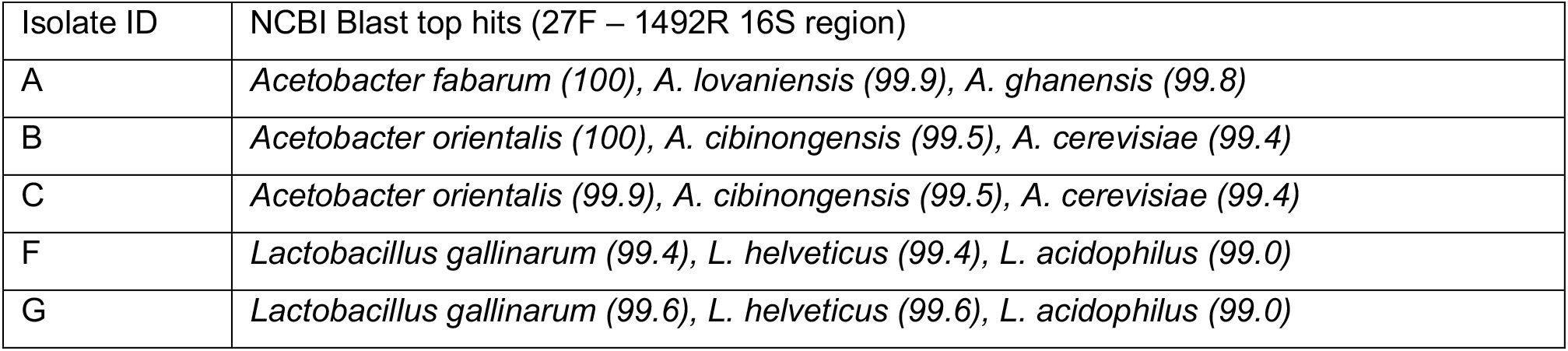
Identity of the five isolates used in synthetic diversity communities with percent identity probabilities in brackets.

| Isolate ID | NCBI Blast top hits (27F – 1492R 16S region) |
| --- | --- |
| A | *Acetobacter fabarum* (100), *A. lovaniensis* (99.9), *A. ghanensis* (99.8) |
| B | *Acetobacter orientalis* (100), *A. cibinongensis* (99.5), *A. cerevisiae* (99.4) |
| C | *Acetobacter orientalis* (99.9), *A. cibinongensis* (99.5), *A. cerevisiae* (99.4) |
| F | *Lactobacillus gallinarum* (99.4), *L. helveticus* (99.4), *L. acidophilus* (99.0) |
| G | *Lactobacillus gallinarum* (99.6), *L. helveticus* (99.6), *L. acidophilus* (99.0) |

**Table S2:** Number of cycles used for PCR amplification in Nanopore sequencing.

| Transfer-community | PCR cycles used |
| --- | --- |
| 1-full | 25 |
| 1-med | 25 |
| 1-low | 23 |
| 1-synt | 25 |
| 5-full | 25 |
| 5-med | 25 |
| 5-low | 24 |
| 5-synt | 30 |
| 17-full | 24 |
| 17-med | 24 |
| 17-low | 24 |
| 17-synt | 24 |

**Table S3:** Separate excel file with details of bioinformatic clustering.

